# Discovery of microbial intergenic features with genomic language modeling and multimodal search

**DOI:** 10.64898/2026.09.15.751765

**Authors:** Nicolo Zulaybar, Matt Tranzillo, Rachel Silverstein, Yunha Hwang, Andre Cornman

## Abstract

Systematic characterization of microbial noncoding regions is limited by two distinct challenges: discovery of conserved sequence features without predefined motifs and functional interpretation of newly identified elements. We address these challenges by training a sparse autoencoder on genomic language model (gLM2) representations to identify intergenic sequence features without prior annotation, and by implementing multimodal search to generate functional hypotheses from conserved associations with neighboring proteins, RNA families, and genomic organization. This framework uncovered divergent, previously uncharacterized noncoding elements, including candidate regulatory DNA sequences and structured RNAs not captured by existing annotation models. gLM2-derived intergenic features can be explored through SeqHub’s multimodal search, freely available for academic use at seqhub.org.

## Main

Microbial genomes and metagenomes contain an enormous diversity of biological sequences, much of which remains functionally unexplored. Large-scale sequence databases have enabled systematic analysis of protein-coding regions, revealing vast numbers of evolutionarily diverse proteins, many of which remain unannotated [1]. By contrast, the noncoding sequences between genes have been studied far less systematically at a large scale.

Microbial noncoding regions encode diverse elements that regulate gene expression or perform molecular functions directly. These include promoters, transcription-factor binding sites, transcriptional terminators, untranslated regulatory elements, riboswitches, small RNAs, and other structured noncoding RNAs [2]. Unlike protein-coding genes, such elements can be short, rapidly evolving, or constrained primarily by secondary structure rather than primary sequence, making them difficult to recognize by conventional sequence similarity searches [3, 4]. Their boundaries are also often unknown, and related elements can differ in length, sequence, and position relative to neighboring genes. Consequently, many functional intergenic elements cannot be easily discovered by searching for predefined motifs.

Many microbial noncoding elements are best understood through their genomic context. Their position relative to neighboring genes, and conservation of these associations across evolution, can reveal both the function of the element and the regulation of nearby genes. Pho boxes, for example, mark PhoB-regulated genes involved in phosphate response [5], while riboswitch function can often be inferred from the metabolic genes they regulate [6]. Clustered Regularly Interspaced Short Palindromic Repeat (CRISPR) -spacer arrays are similarly linked to adjacent Cas proteins, connecting distinctive noncoding architectures to RNA-guided immunity [7]. More recently, bridge RNAs were identified within IS110-family elements, where their association with cognate recombinases helped reveal an RNA-guided mechanism of DNA recombination [8]. Together, these examples illustrate how genomic context can provide functional information that is not apparent from the noncoding sequence alone.

Systematically exploiting these relationships requires two capabilities not well supported by existing genomic analysis methods. First, de novo discovery differs fundamentally from sequence search because the relevant feature is not known in advance: there may be no predefined query, motif length, sequence pattern, or structural architecture to search for. Sequence- and covariance-based methods are powerful once a query or candidate family has been defined [3, 9, 10, 11], but the initial discovery problem requires identifying recurrent signals directly from genomic sequence. Second, even after a sequence feature is discovered, its biological role may remain unclear. Conserved associations with neighboring genes, protein families, RNA elements and genomic organization can provide functional clues, but systematically identifying these relationships across large genome databases is difficult because these modalities are not jointly searchable.

Here, we combine genomic language modeling [12] with multimodal search to enable context-aware discovery of microbial intergenic features. We use gLM2 [13], a genomic language model trained on microbial genome and metagenome sequences, to identify conserved sequence features in their native genomic context without requiring a predefined motif or annotated family. We then use SeqHub [14], a multimodal microbial genomic context search software freely available for academic use, to search across protein sequences, protein domains, known RNA families, genomic neighborhoods, and gLM2-derived intergenic sequence features to characterize the genomic associations of candidate elements. Together, these approaches connect pattern discovery with contextual interpretation, allowing intergenic features to be identified and traced across genomes according to the conserved genes and genomic features with which they associate. We apply this framework to identify previously uncharacterized intergenic elements associated with conserved microbial functions, providing a general approach for interrogating the noncoding fraction of microbial genomes at database scale.

To obtain interpretable features from gLM2, we trained a sparse autoencoder (SAE) [15] on layer-24 activations from genomic context windows containing interleaved protein-coding and intergenic sequence (Fig. 1a,b). The SAE contained 16,384 latent features and was trained on 10^9^ intergenic tokens, with the objective restricted to nucleotide positions. We then applied the frozen gLM2-SAE model across the OpenGenome database [13] comprising 131,744 microbial genomes, ∼400 million protein-coding genes and ∼300 million intergenic regions. Feature spans were defined by grouping firing positions separated by *≤* 3 nt and discarding spans *<* 8 nt. For downstream analyses, we retained 3,656 features observed in at least 50 contigs.

**Figure 1.**
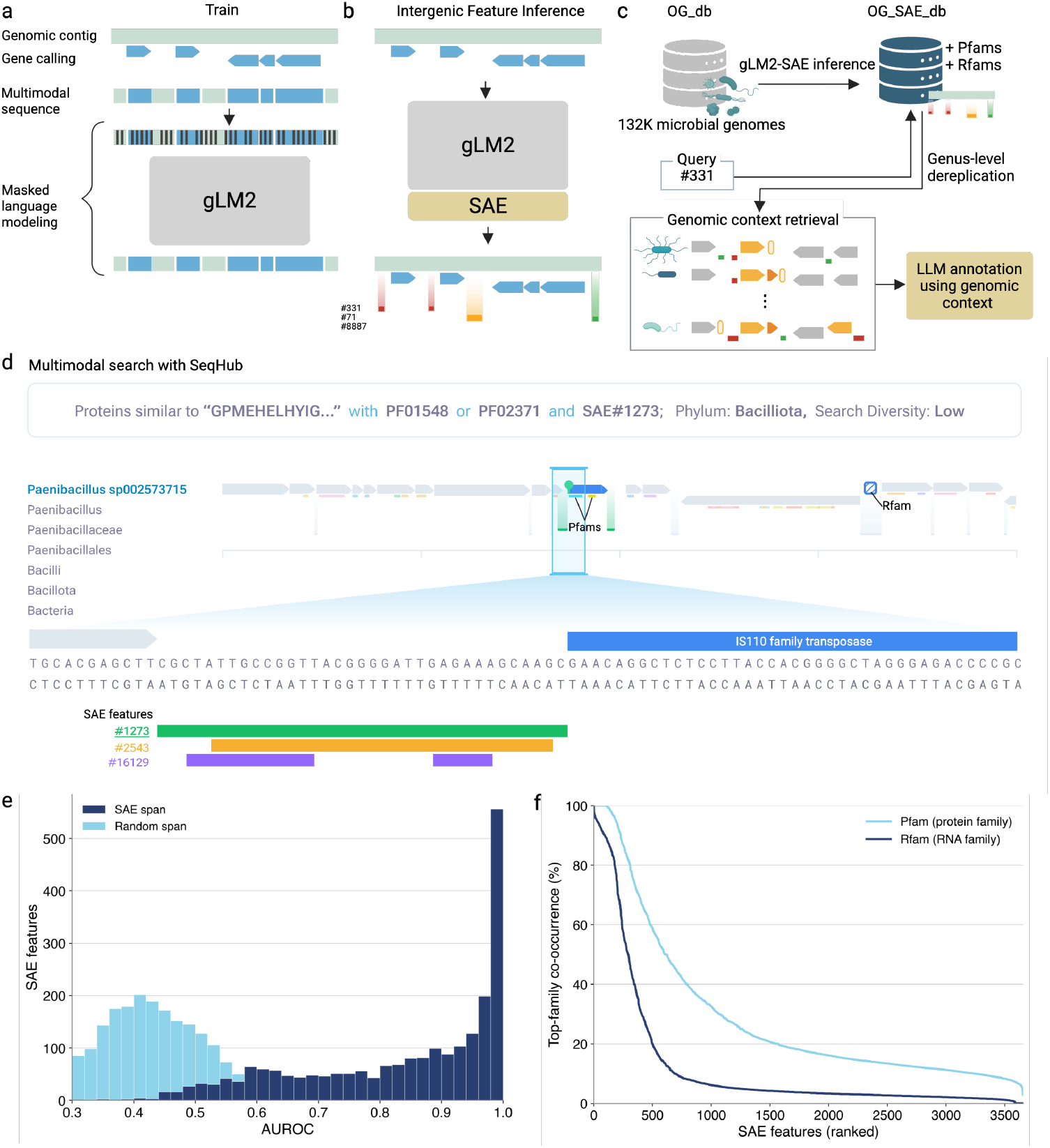
Discovery and contextual annotation of microbial intergenic features with gLM2 and SeqHub. **(A)** gLM2 is trained on multimodal genomic sequences containing interleaved proteincoding and intergenic regions using masked language modeling. **(B)** A sparse autoencoder (SAE) trained on gLM2 intergenic embeddings identifies interpretable nucleotide-level features; nearby activating positions are grouped into genomic feature spans. **(C)** For each SAE feature, high-activating loci are retrieved across diverse genomes and annotated using neighboring genes, Pfam and Rfam families, taxonomy and gene organization. These evolutionarily conserved genomic contexts are summarized to generate putative feature annotations. **(D)** SeqHub enables multimodal search across protein similarity, Pfam and Rfam families, SAE features and taxonomy to retrieve conserved genomic contexts. **(E)** SAE features capture sequencespecific motifs that generalize across genera. For each feature, the 150 highest-activating intergenic regions per genus were paired with length-matched negative regions from the same genomic segments and split 80:20 into training and held-out sets. STREME motifs learned from the training positives were evaluated on held-out pairs. SAE-associated regions yielded a median AUROC of 0.893 (blue), compared with 0.429 when positive regions were replaced by random intergenic regions from the same contigs (light blue). **(F)** Distribution of genomic-context associations across SAE features. For each feature, Pfam and Rfam co-occurrence were defined as the fraction of genus-dereplicated sampled contigs containing its most frequently associated family. Features are ranked independently by this co-occurrence fraction for Pfam and Rfam; they-axis shows the corresponding top-family co-occurrence percentage.

We next constructed an annotation pipeline to characterize the genomic patterns captured by individual SAE features (Fig. 1c). For each feature, we retrieved the genomic contexts surrounding its 100 highest-activating spans and dereplicated these results at the genus level to limit signals dominated by closely related genomes. Each context was annotated with Pfam protein-family annotations [16], Rfam RNA-family annotations [17], functional predictions [18], gene orientation and position, and taxonomic information. These contextual summaries were then provided to a large language model to generate a putative description of the recurrent sequence pattern and its genomic associations. Feature annotations were therefore based not on SAE activation alone, but on the evolutionary distribution of each feature and the genes, protein domains and known RNA elements that repeatedly co-occurred with it across diverse genomes.

To enable analysis of SAE features with conserved genomic context, we integrated the resulting SAE features into SeqHub and implemented a multimodal genomic-context search system (Fig. 1d). Queries can combine one or more protein sequences, SAE feature identifiers, Pfam families and Rfam families using AND, OR and NOT operators, and can be constrained by taxonomic scope and diversity level. This makes it possible to begin the discovery process with either a candidate intergenic pattern or a neighboring biological feature and iteratively identify the conserved genomic architectures that connect them.

To assess whether SAE firing spans capture reproducible primary-sequence signals, for each feature we selected its 150 highest-activating intergenic regions, at most one per genus, and paired each with a length-matched negative drawn from an intergenic region in the same genomic segment (see methods). We used STREME [19] to discover motifs from the training positives against their matched negatives and scored its top-ranked motif on the held-out sequences. The median held-out AUROC was 0.893, and 48% of the tested features exceeded an Area Under the Receiver Operating Characteristic curve (AUROC) of 0.9 (Fig. 1e). To determine whether this performance could instead arise from genome- or contig-specific sequence composition, we repeated the analysis using positive regions sampled uniformly from other intergenic regions on the same contigs. In this control, the median AUROC was 0.429, compared with 0.893 for SAE-associated regions, and in 96% of features the control scored lower (Wilcoxon signed-rank test, *P <* 10^*−*15^). These results indicate that a substantial fraction of gLM2-derived intergenic features capture sequence-specific signals that generalize across genera.

Because the function of an intergenic element can often be inferred from the genes with which it consistently co-occurs, we next examined the genomic associations of SAE features. Across genus-level dereplicated contigs, 605 features (16.5%) co-occurred with the same Pfam domain on more than half of associated contigs, and 290 (7.9%) showed the same level of association with an Rfam (Fig. 1f). These conserved associations suggest that many gLM2 intergenic features are embedded within specific biological systems, providing a route to infer their potential function from genomic context.

We next examined representative SAE features associated with distinct classes of intergenic elements, including putative operator sequences, divergent members of known RNA families, and previously uncharacterized structured RNAs (Fig. 2). Feature #790 consistently fired in an AT-rich region upstream of a conserved YoaS-YozG operon across diverse phyla, encoding DUF2975 and a Cro/C1-like HTH repressor, respectively. The activated regions contained sequence-diverse inverted repeats, consistent with putative operator sites for autoregulation by the adjacent HTH repressor [20] (Fig. 2a). Feature #7917 was associated with bacterial Selenocysteine Insertion Sequence (SECIS) elements; among its 100 highest-activating regions, only 9 overlapped the SECIS-3 (RF01989) and SECIS-4 (RF01990) Rfam families. Many of the remaining loci nevertheless showed the characteristic organization of bacterial selenoprotein genes, with an in-frame UGA codon followed by a nearby SECIS element [21]. Some of the associated proteins included well characterized selenoproteins such as formate dehydrogenase *α* subunits and other molybdopterin oxidoreductases [22], while many consisted of less-characterized proteins with selenocysteine predicted to be incorporated at conserved catalytic Cys/Sec motifs [23] (Fig. 2b). Features #14380 and #6752 were similarly associated with metal-responsive riboswitches: 15 of the 100 highest-activating #14380 regions overlapped the M-box magnesium riboswitch (RF00380), whereas 21 and 6 of the top #6752 regions overlapped the yybP-ykoY manganese riboswitch (RF00080) and TerC putative riboswitch (RF03067) respectively [24, 25, 26, 27]. For highly activating regions without an Infernal hit, Se-qHub searches combining the SAE feature with downstream membrane-transporter context enriched homologous loci, from which CMfinder [11] and R-scape [28] identified RNA structures supported by significant covariation and distinct from the corresponding Rfam models. (Fig. 2c). Finally, feature #13341 marked an unannotated RNA element conserved near ribosomal proteins uS9 and uS2 and frequently associated with elongation factor Ts (EF-Ts) and uL13, with the strongest representation in the Bacteroidota phylum. A multimodal search combining uS2 with feature #13341 identified homologous regions that yielded a conserved covarying secondary structure with no Rfam match. Its conserved association with ribosomal-protein loci suggests a regulatory role, similar to autogenous feedback regulation by structured RNA in bacterial ribosomal-protein operons [29] (Fig. 2d). Together, these results show that coupling genomic language model features with multi-modal context search provides a general strategy to systematically uncover and interpret microbial intergenic elements beyond the reach of existing annotation frameworks.

**Figure 2.**
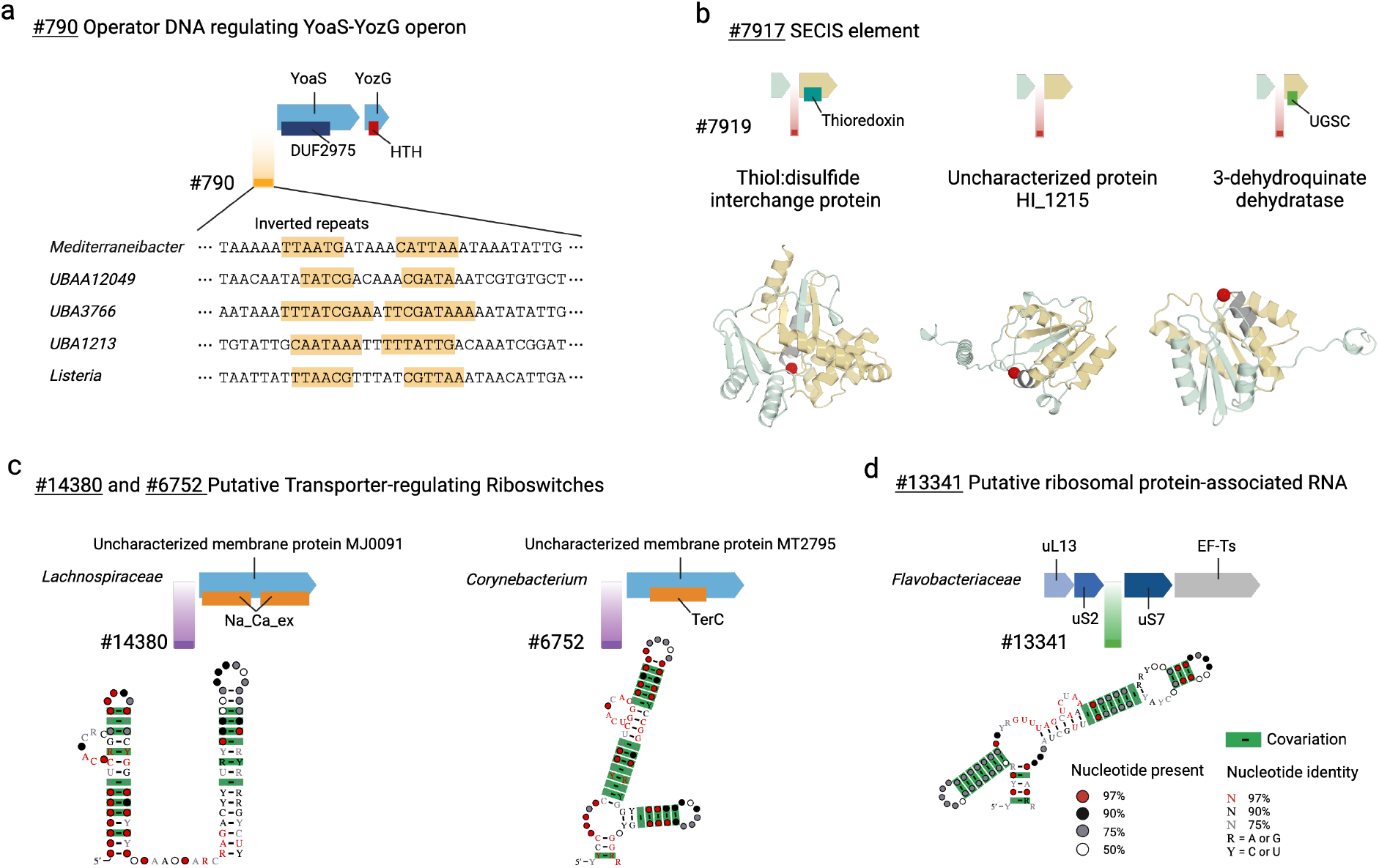
Representative intergenic elements identified by gLM2 features and contextualized with SeqHub. **(A)** Feature #790 marks an AT-rich region upstream of the conserved YoaS-YozG operon in Bacillota. Representative activating sequences contain sequence-diverse inverted repeats, consistent with putative operator sites for the adjacent Cro/C1-like HTH repressor. **(B)** Bacterial SECIS elements identified by feature #7917 that are not annotated by known Rfams using Infernal. Representative examples include established and less-characterized selenoproteins; predicted selenocysteine positions are highlighted in red in the corresponding protein structures. **(C)** Features #14380 and #6752 identify putative riboswitches of uncharacterized membrane transporters. SeqHub searches combining each SAE feature with downstream membrane-transporter context recovered divergent loci lacking Infernal hits; covariation analysis identified conserved, significantly covarying RNA structures. **(D)** Feature #13341 identifies an unannotated RNA element conserved near ribosomal protein loci, including small ribosomal subunit S2 (uS2), small ribosomal subunit S9 (uS9), Elongation Factor Ts (EF-Ts) and large ribosomal subunit L13 (uL13). Multimodal retrieval followed by covariation analysis identified a conserved secondary structure with no Rfam match. CMfinder output scores are found in Supplemental Table S1.

## Methods

### Sparse autoencoder training

We trained a sparse autoencoder (SAE) on residual-stream activations from gLM2-650M, a 33-layer bidirectional mixed-modality genomic language model with a hidden dimension of 1,280. gLM2 represents coding sequences as amino acids and intergenic sequences as nucleotides, interleaved in genomic order (Fig. 1A). Training sequences were sampled from the OG dataset. To reduce redundancy from over-represented isolate genomes, 4,096-token windows were centered on 50%-sequence-identity cluster representatives (at 70% coverage) determined by MMseqs2 [30]. We extracted residual-stream activations after transformer block 24 with the language model frozen. Complete genomic windows were passed through gLM2, but only nucleotide positions were retained for SAE training, so that each training position retained the context of neighboring genes while the reconstruction objective was restricted to intergenic sequence.

The SAE used a BatchTopK architecture with 16,384 latent features and *K* = 8, retaining on average eight active features per position [31]. The loss comprised reconstruction mean-squared error and an auxiliary reconstruction loss for features inactive for 300 optimization steps. The decoder bias was initialized to the geometric median of the activation distribution to improve feature utilization [32]. We trained the SAE on 10^9^ intergenic positions using Adam (*β*_1_ = 0, *β*_2_ = 0.95), a learning rate of 7 *×* 10^*−*5^, 1,000 warm-up steps and a global batch size of 32,768 across eight H100 GPUs. Training code was adapted from SAELens [33].

### Multimodal genomic-context search

To make SAE features searchable across a large genomic database, we applied the frozen gLM2-SAE model to all contig segments in OG and recorded feature activations at each intergenic position. Firing positions for a given feature were grouped into spans when separated by no more than three nucleotides. Spans shorter than eight nucleotides were discarded to exclude isolated or sporadic activations, and the remaining spans were stored with their genomic coordinates and activation values. For downstream analyses, we retained 3,656 features with spans detected on at least 50 contigs. The distributions of median span length, prevalence across OG contigs and number of features per intergenic region are shown in Supplementary Figs. S1–S3.

We integrated these spans into SeqHub together with protein sequences, Pfam and Rfam annotations, gene coordinates and taxonomy, linked through shared contig and gene identifiers. Queries can combine protein sequences, SAE feature identifiers and Pfam or Rfam accessions using AND, OR and NOT operators and can be restricted by taxonomy. Protein queries are resolved by embedding similarity against protein-coding genes, whereas SAE features and Pfam and Rfam families are matched directly to their indexed occurrences. Protein searches can additionally be restricted to cluster representatives to reduce sequence redundancy at 70%, 50% and 30% sequence identity.

### Primary-sequence motif evaluation

We tested whether SAE features captured reproducible primary-sequence motifs using STREME (MEME Suite v5.5.7) [19]. Features were considered if they fired on at least 2,000 but no more than 25% of OG contigs. For each feature, we selected its highest-activating intergenic region from each of up to 150 genera, retaining at most one region per genus. Each positive region was paired with a length-matched intergenic region from the same contig in which the feature did not fire. Features yielding at least 150 matched positive–negative pairs were evaluated, resulting in 2,234 features.

Matched pairs were divided 80:20 into training and test sets, with both members of each pair assigned to the same set. Because each pair represented a distinct genus, genera did not overlap between the training and test sets. STREME was run on the training positives using the matched training negatives as controls. The top-ranked motif was converted to a log-odds position-weight matrix using the nucleotide composition of the training negatives. Each held-out sequence was assigned its maximum motif score, and motif performance was measured by the area under the receiver operating characteristic curve (AUROC) across the held-out positives and negatives.

As a control, we repeated the analysis after replacing each positive with an intergenic region sampled from the same contig, leaving the contig set, genus dereplication, negatives and train–test split unchanged. Paired AUROCs were compared using a one-sided Wilcoxon signed-rank test.

### Genomic context and co-occurrence analysis

For each SAE feature, we retained the highest-activating contig from each of up to 1,000 distinct genera. The genomic neighborhood was defined as the five annotated genes upstream and downstream of the intergenic region containing the span.

Pfam domains were identified using pyhmmer (v0.10.14) hmmsearch [34] against Pfam-A v38.0 [35] at each family’s curated gathering threshold, and counted once per gene irrespective of the number of domain occurrences. Rfams were annotated on intergenic sequence using nhmmer (*E ≤* 10^*−*3^) [9] on Rfam version 15.1 [17]; nhmmer was used for database-scale annotation because Infernal was computationally prohibitive at this scale.

For each feature, Pfam and Rfam co-occurrence were defined as the fraction of sampled contigs containing its most frequently associated family. Analyses were restricted to the 3,656 features detected on at least 50 contigs.

## LLM-based feature annotation

To assign putative functions to SAE features, we summarized their genomic contexts and interpreted these summaries with a large language model. For each feature, contigs were ranked by peak activation and genus-dereplicated to retain up to 20 sites from distinct genera, examining up to 2,000 candidates per feature.

For each site, we recorded the firing-span sequence with 30 bp of flanking sequence, its position and orientation relative to neighboring genes, and overlap with Rfam annotations. Feature-level summaries additionally included neighboring Pfams and Rfams, genomic prevalence, and taxonomic distribution across genera and phyla. These summaries were provided to Claude Opus 5, which returned a feature name, functional description, supporting evidence, and a confidence level. These summaries are available at https://seqhub.org.

## RNA structural covariation analysis

We first examined the features’ overlap with known Rfam annotations, which was defined as the union of two searches over each feature’s 100 highest-activating, genus-dereplicated regions: the database-scale nhmmer annotation above, and a more sensitive covariance-model search of the sequence surrounding each region’s highest-activating firing run (Infernal v1.1.5 cmscan, Rfam gathering thresholds). A region was counted once per family, and only when the hit overlapped that firing run.

To test whether SAE features marked conserved RNA secondary structures, we analyzed intergenic regions in which a feature co-occurred with homologs of a specified query protein. Protein-anchored sampling restricted each analysis to a consistent genomic context. For each query consisting of a protein-feature pair, we retrieved 60 co-search matches for each flank of the protein and counted the loci containing a feature firing span in the corresponding intergenic region. We retained the flank with more firing loci and excluded pairs for which the feature did not fire on either flank.

We then retrieved up to 1,000 loci in which homologs of the query protein co-occurred with the SAE feature. From each locus, we extracted the complete intergenic region on the selected flank. Sequences were oriented relative to the matched gene, such that upstream sequences ended at its start codon, and were supplied to CMfinder [11] without prior alignment. Candidate structures were inferred using CMfinder v0.4.1.9 from the 100 loci whose protein homologs were most similar to the query. Covariation in each resulting alignment was evaluated using R-scape v2.0.4a [28] at E-value *≤* 0.05. For alignments containing at least one significantly covarying base pair, we built and calibrated a covariance model using Infernal v1.1.5 (cmbuild and cmcalibrate) [3] and searched it against all extracted sequences using cmsearch at E-value *≤* 10^*−*3^. We report the proportion of extracted sequences recovered by each model and the corresponding motifs scores reported by CMfinder’s ScoreMotif.pl (RNAphylo scores and hmmpair ranks) in Supplemental Table S1. Secondary structures were visualized using R2R [36].

## Acknowledgements

We thank SeqHub users for feedback and suggestions that helped shape the intergenic analysis, and Alex Bateman and Jordan Hoff for insightful discussions. Employees at Tatta Bio are supported by Schmidt Futures through Grant G-24-67500 and by Gordon and Betty Moore Foundation through Grant #13344 to Tatta Bio.

## Author Contributions

Conceptualization: YH, AC

Methodology: YH, RS, AC

Investigation: MT, NZ, YH, RS, AC

Data curation: YH, AC

Validation: MT, NZ, RS, YH, AC

Formal analysis: YH, AC

Resource: YH, AC

Software: MT, NZ, AC

Project Administration: YH, AC

Visualization: MT, NZ, YH, AC

Funding acquisition: YH, AC

Supervision: YH, AC

Writing-original draft: YH, AC

Writing-review & editing: MT, NZ, RS, YH, AC

## Data and code availability

The OpenGenome (OG) dataset used for SAE training is available on HuggingFace at https://huggingface.co/datasets/tattabio/OG. The gLM2 SAE model is available at https://huggingface.co/tattabio/gLM2_650M_sae. Inference code is available at https://github.com/TattaBio/gLM2_SAE. Multimodal search, recorded feature spans, automated annotations of features across the OG database are available on SeqHub, a free software for academic and noncommercial use.

## Supplementary Material

**Figure S1:**
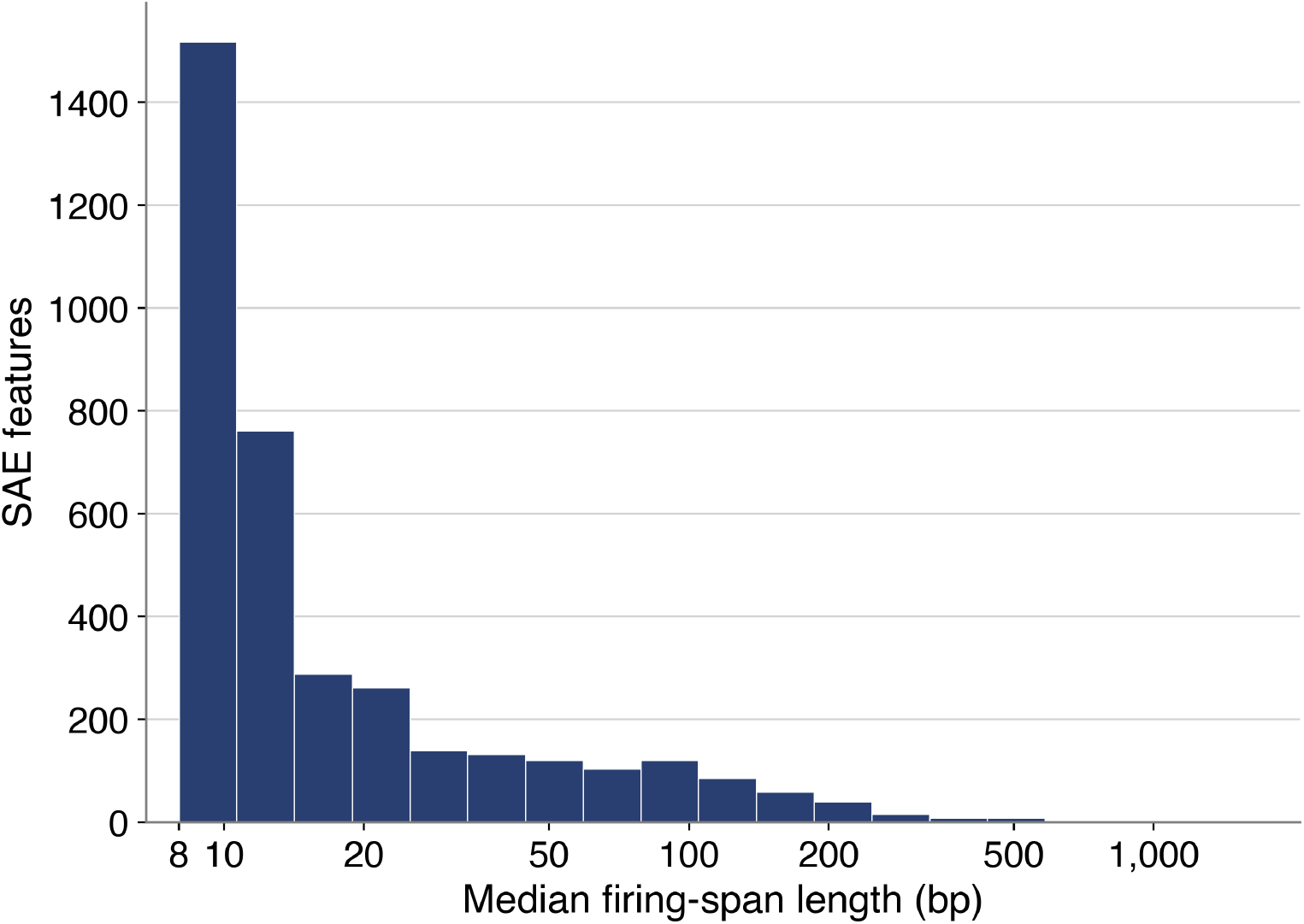
Firing-span lengths of SAE features. Median length of a feature’s firing spans for the 3,656 features firing on at least 50 contigs. Spans shorter than 8 bp are discarded during feature extraction. The x axis is logarithmically scaled.

**Figure S2:**
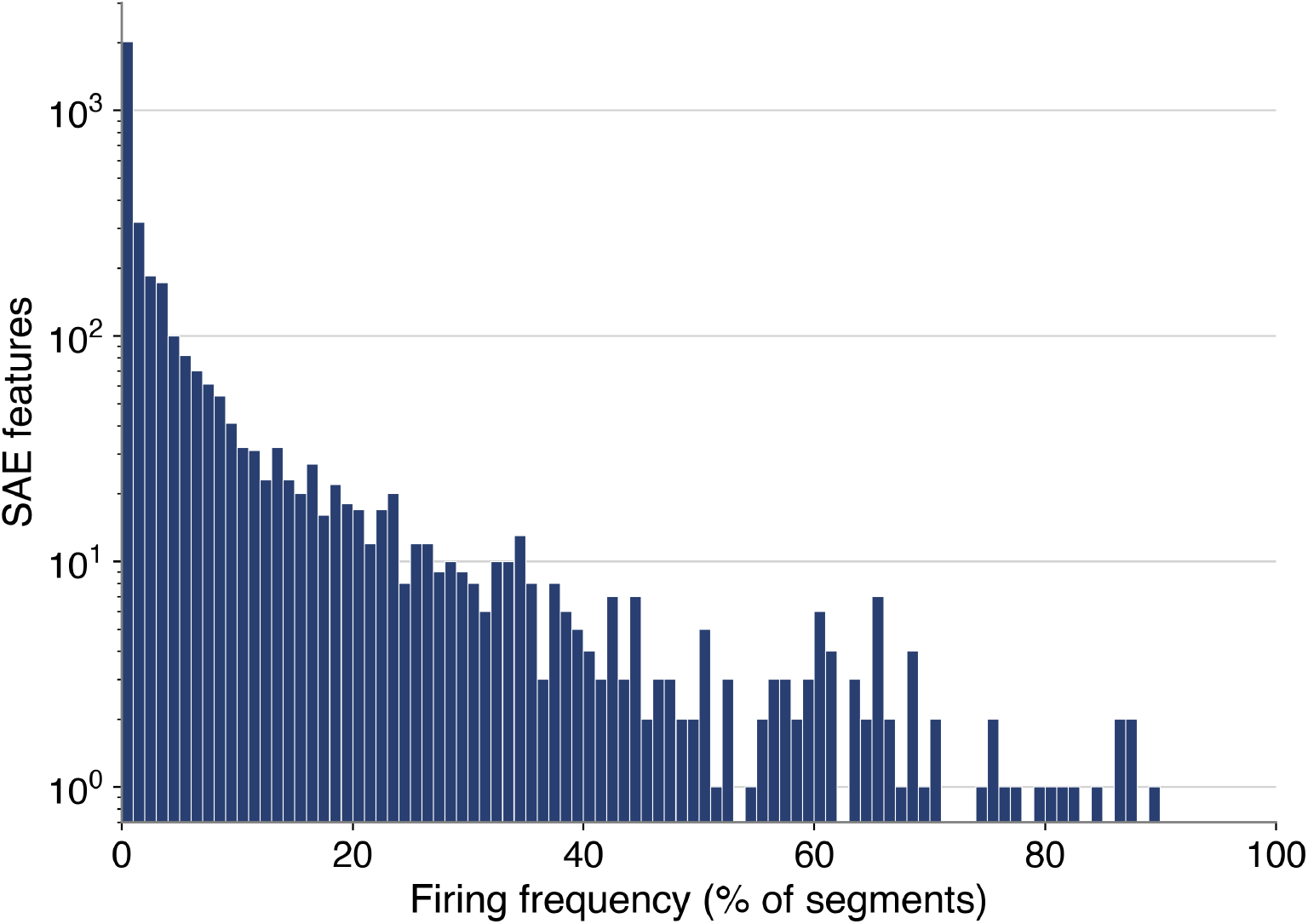
Prevalence of SAE features across OpenGenome contigs. Distribution of the percentage of OpenGenome contigs containing at least one firing span for each SAE feature. The y axis shows the number of features on a logarithmic scale.

**Figure S3:**
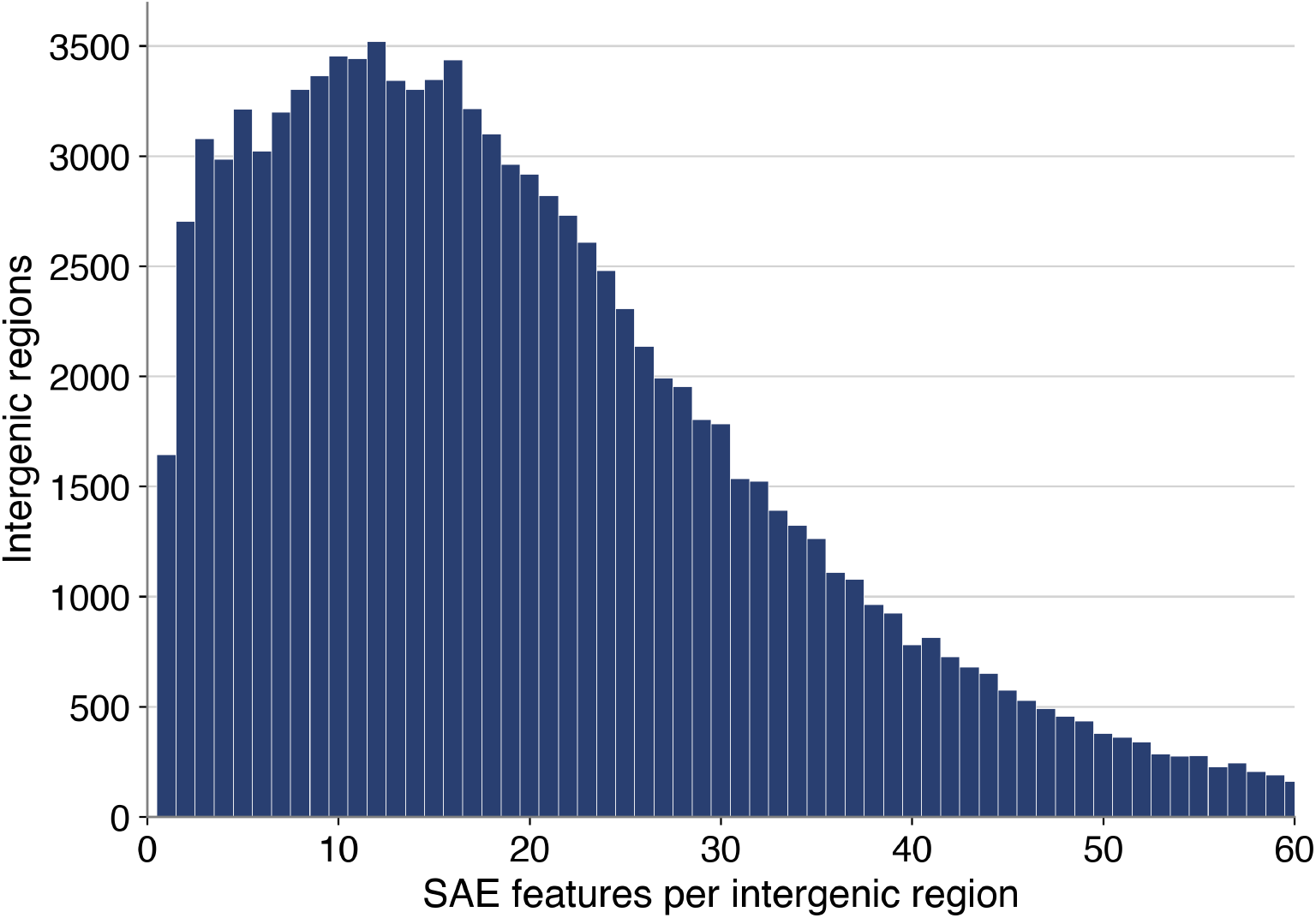
Number of SAE features per intergenic region. Distribution of the number of distinct SAE features with a firing span in each intergenic region from a random sample of 2,000 OpenGenome contigs.

**Table S1:** CMfinder motifs recovered from SAE-guided co-searches. For each protein anchor, SeqHub was queried with the protein sequence AND the SAE feature, and the full intergenic gap flanking each retrieved gene was extracted; CMfinder was run on the first 100 such sequences and every proposed motif was tested with R-scape. The single best-supported motif per locus is shown. *N*_*seq*_ is the number of sequences in that motif’s alignment, out of the 100 given to CMfinder; CMfinder keeps only the subset in which it can place the motif. *Obs*.*/exp*. are R-scape’s observed and expected numbers of significantly covarying base pairs at *E ≤* 0.05. The expected value is the number detectable given the alignment’s phylogenetic depth. A ratio near 1 indicates covariation at the level a genuine conserved structure of that depth and diversity would produce. *Self-verified* is the number of distinct sequences, out of all extracted for that locus, that the calibrated covariance model re-detects by cmsearch at *E ≤* 10^*−*3^. *hmmpair rank* is the CMfinder `ScoreMotif.pl hmmpairRank`, the motif’s rank against a reference set of *∼*269,000 motifs, where a lower rank is better. *Motif score* is the combined score output from `ScoreMotif.pl`.

| Locus (protein anchor) | $N_{seq}$ | Obs./exp. | Self-verified | hmmpair rank | Motif score |
| --- | --- | --- | --- | --- | --- |
| uS2 / feature #13341 | 97 | 19 / 19.2 | 213/586 (36%) | 37,277 | −222 |
| MT2795 / feature #6752 | 96 | 18 / 17.7 | 628/825 (76%) | 9,874 | −99 |
| MJ0091 / feature #14380 | 45 | 14 / 14.0 | 45/62 (73%) | 478 | −418 |

## References

Georgios A Pavlopoulos, Fotis A Baltoumas, Sirui Liu, Oguz Selvitopi, Antonio Pedro Camargo, Stephen Nayfach, Ariful Azad, Simon Roux, Lee Call, Natalia N Ivanova, I Min Chen, David Paez-Espino, Evangelos Karatzas, Novel Metagenome Protein Families Consortium, Ioannis Iliopoulos, Konstantinos Konstantinidis, James M Tiedje, Jennifer Pett-Ridge, David Baker, Axel Visel, Christos A Ouzounis, Sergey Ovchinnikov, Aydin Buluç, and Nikos C Kyrpides. Unraveling the functional dark matter through global metagenomics. Nature, 622(7983): 594–602, October 2023.

Dmitry A Rodionov. Comparative genomic reconstruction of transcriptional regulatory networks in bacteria. Chem. Rev., 107(8):3467–3497, August 2007.

Eric P Nawrocki and Sean R Eddy. Infernal 1.1: 100-fold faster RNA homology searches. Bioinformatics, 29(22):2933–2935, November 2013.

Zasha Weinberg, Joy X Wang, Jarrod Bogue, Jingying Yang, Keith Corbino, Ryan H Moy, and Ronald R Breaker. Comparative genomics reveals 104 candidate structured RNAs from bacteria, archaea, and their metagenomes. Genome Biol., 11(3):R31, March 2010.

Kozo Makino, Hideo Shinagawa, Mitsuko Amemura, Sigenobu Kimura, Atsuo Nakata, and Akira Ishihama. Regulation of the phosphate regulon of escherichia coli. J. Mol. Biol., 203(1): 85–95, September 1988.

Madeline E Sherlock and Ronald R Breaker. Former orphan riboswitches reveal unexplored areas of bacterial metabolism, signaling, and gene control processes. RNA, 26(6):675–693, June 2020.

Ruud Jansen, Jan D A van Embden, Wim Gaastra, and Leo M Schouls. Identification of genes that are associated with DNA repeats in prokaryotes. Mol. Microbiol., 43(6):1565–1575, March 2002.

Matthew G Durrant, Nicholas T Perry, James J Pai, Aditya R Jangid, Januka S Athukoralage, Masahiro Hiraizumi, John P McSpedon, April Pawluk, Hiroshi Nishimasu, Silvana Konermann, and Patrick D Hsu. Bridge RNAs direct programmable recombination of target and donor DNA. Nature, 630(8018):984–993, June 2024.

Travis J Wheeler and Sean R Eddy. nhmmer: DNA homology search with profile HMMs. Bioinformatics, 29(19):2487–2489, October 2013.

Sukhwan Park, Kieran Didi, Andrew Favor, Anton Bushuiev, Soohyun Kim, Milot Mirdita, and Martin Steinegger. Fast remote nucleotide sequence alignment with riboseek. bioRxiv, page 2026.07.31.741718, July 2026.

Zizhen Yao, Zasha Weinberg, and Walter L Ruzzo. CMfinder–a covariance model based RNA motif finding algorithm. Bioinformatics, 22(4):445–452, February 2006.

Yunha Hwang, Andre L Cornman, Elizabeth H Kellogg, Sergey Ovchinnikov, and Peter R Girguis. Genomic language model predicts protein co-regulation and function. Nat. Commun., 15(1):2880, April 2024.

Andre Cornman, Jacob West-Roberts, Antonio Pedro Camargo, Simon Roux, Martin Bera-cochea, Milot Mirdita, Sergey Ovchinnikov, and Yunha Hwang. The OMG dataset: An open MetaGenomic corpus for mixed-modality genomic language modeling. In The Thirteenth International Conference on Learning Representations, October 2024.

Nishant Jha, Joshua Kravitz, Jacob West-Roberts, Antonio Camargo, Simon Roux, Andre Cornman, and Yunha Hwang. Gaia: A context-aware sequence search and discovery tool for microbial proteins. bioRxiv, page 2024.11.19.624387, November 2024.

Hoagy Cunningham, Aidan Ewart, Logan Riggs, Robert Huben, and Lee Sharkey. Sparse autoencoders find highly interpretable features in language models. arXiv [cs.LG], September 2023.

Jaina Mistry, Sara Chuguransky, Lowri Williams, Matloob Qureshi, Gustavo A Salazar, Erik L L Sonnhammer, Silvio C E Tosatto, Lisanna Paladin, Shriya Raj, Lorna J Richardson, Robert D Finn, and Alex Bateman. Pfam: The protein families database in 2021. Nucleic Acids Res., 49(D1):D412–D419, January 2021.

Nancy Ontiveros-Palacios, Emma Cooke, Eric P Nawrocki, Sandra Triebel, Manja Marz, Elena Rivas, Sam Griffiths-Jones, Anton I Petrov, Alex Bateman, and Blake Sweeney. Rfam 15: RNA families database in 2025. Nucleic Acids Res., 53(D1):D258–D267, January 2025.

Nishant Jha, Joshua Kravitz, Jacob West-Roberts, Cong Lu, Antonio Pedro Camargo, Simon Roux, Andre Cornman, and Yunha Hwang. Gaia: An AI-enabled genomic context-aware platform for protein sequence annotation. Sci. Adv., 11(25):eadv5109, June 2025.

Timothy L Bailey. STREME: accurate and versatile sequence motif discovery. Bioinformatics, 37(18):2834–2840, September 2021.

A Johnson, B J Meyer, and M Ptashne. Mechanism of action of the cro protein of bacteriophage lambda. Proc. Natl. Acad. Sci. U. S. A., 75(4):1783–1787, April 1978.

A Hüttenhofer, E Westhof, and A Böck. Solution structure of mRNA hairpins promoting selenocysteine incorporation in escherichia coli and their base-specific interaction with special elongation factor SELB. RNA, 2(4):354–366, April 1996.

Yan Zhang, Steffen Rump, and Vadim N Gladyshev. Comparative genomics and evolution of molybdenum utilization. Coord. Chem. Rev., 255(9-10):1206–1217, May 2011.

Valentina A Shchedrina, Sergey V Novoselov, Mikalai Yu Malinouski, and Vadim N Gladyshev. Identification and characterization of a selenoprotein family containing a diselenide bond in a redox motif. Proc. Natl. Acad. Sci. U. S. A., 104(35):13919–13924, August 2007.

L Argaman, R Hershberg, J Vogel, G Bejerano, E G Wagner, H Margalit, and S Altuvia. Novel small RNA-encoding genes in the intergenic regions of escherichia coli. Curr. Biol., 11 (12):941–950, June 2001.

Michael Dambach, Melissa Sandoval, Taylor B Updegrove, Vivek Anantharaman, L Aravind, Lauren S Waters, and Gisela Storz. The ubiquitous yybP-ykoY riboswitch is a manganese-responsive regulatory element. Mol. Cell, 57(6):1099–1109, March 2015.

Jeffrey E Barrick, Keith A Corbino, Wade C Winkler, Ali Nahvi, Maumita Mandal, Jennifer Collins, Mark Lee, Adam Roth, Narasimhan Sudarsan, Inbal Jona, J Kenneth Wickiser, and Ronald R Breaker. New RNA motifs suggest an expanded scope for riboswitches in bacterial genetic control. Proc. Natl. Acad. Sci. U. S. A., 101(17):6421–6426, April 2004.

Charles E Dann, 3rd, Catherine A Wakeman, Cecelia L Sieling, Stephanie C Baker, Irnov Irnov, and Wade C Winkler. Structure and mechanism of a metal-sensing regulatory RNA. Cell, 130(5):878–892, September 2007.

Elena Rivas, Jody Clements, and Sean R Eddy. Estimating the power of sequence covariation for detecting conserved RNA structure. Bioinformatics, 36(10):3072–3076, May 2020.

Leonid V Aseev, Alexandrina A Levandovskaya, Ludmila S Tchufistova, Nadezda V Scaptsova, and Irina V Boni. A new regulatory circuit in ribosomal protein operons: S2-mediated control of the rpsB-tsf expression in vivo. RNA, 14(9):1882–1894, September 2008.

Martin Steinegger and Johannes Söding. MMseqs2 enables sensitive protein sequence searching for the analysis of massive data sets. Nat. Biotechnol., 35(11):1026–1028, November 2017.

Bart Bussmann, Patrick Leask, and Neel Nanda. BatchTopK sparse autoencoders. arXiv [cs.LG], December 2024.

Elana Simon, Etowah Adams, and James Zou. On the relationship between activation outliers and feature death in sparse autoencoders. arXiv [cs.LG], May 2026.

Joseph Bloom, Curt Tigges, Anthony Duong, and David Chanin. Saelens. https://github.com/decoderesearch/SAELens, 2024.

L Steven Johnson, Sean R Eddy, and Elon Portugaly. Hidden markov model speed heuristic and iterative HMM search procedure. BMC Bioinformatics, 11(1):431, August 2010.

Typhaine Paysan-Lafosse, Antonina Andreeva, Matthias Blum, Sara Rocio Chuguransky, Tiago Grego, Beatriz Lazaro Pinto, Gustavo A Salazar, Maxwell L Bileschi, Felipe Llinares-López, Laetitia Meng-Papaxanthos, Lucy J Colwell, Nick V Grishin, R Dustin Schaeffer, Damiano Clementel, Silvio C E Tosatto, Erik Sonnhammer, Valerie Wood, and Alex Bateman. The pfam protein families database: embracing AI/ML. Nucleic Acids Res., 53(D1):D523–D534, January 2025.

Zasha Weinberg and Ronald R Breaker. R2R–software to speed the depiction of aesthetic consensus RNA secondary structures. BMC Bioinformatics, 12(1):3, January 2011.

